# Exon skipping via chimeric antisense U1 snRNAs to correct Retinitis Pigmentosa GTPase-Regulator (RPGR) splice defect

**DOI:** 10.1101/2021.06.26.449721

**Authors:** Giuseppina Covello, Gehan H. Ibrahim, Niccolò Bacchi, Simona Casarosa, Michela Alessandra Denti

**Affiliations:** RNA Biology and Biotechnology Laboratory, Department of Cellular, Computational and Integrative Biology - CIBIO, University of Trento, Trento, Italy; Department of Medical Biochemistry, Faculty of Medicine, Suez Canal University, Round Road, Ismailia, Egypt; Neural Development and Regeneration Laboratory, Department of Cellular, Computational and Integrative Biology - CIBIO, University of Trento, Trento, Italy; CNR Neuroscience Institute, Pisa, Italy; Department of Biology, University of Padova, Padova, Italy

**Keywords:** alternative splicing, U1 snRNA, exon skipping, Retinitis Pigmentosa GTPase-regulator (RPGR), RNA Therapeutics

## Abstract

Inherited retinal dystrophies are caused by mutations in more than 250 genes, each of them carrying several types of mutations that can lead to different clinical phenotypes. Mutations in *Retinitis Pigmentosa GTPase-Regulator* (RPGR) cause X-linked Retinitis pigmentosa (RP). A nucleotide substitution in intron 9 of *RPGR* causes the increase of an alternatively spliced isoform of the mature mRNA, bearing exon 9a (E9a). This introduces a stop codon, leading to truncation of the protein. Aiming at restoring impaired gene expression, we developed an antisense RNA-based therapeutic approach for the skipping of RPGR E9a. We designed a set of specific U1 antisense snRNAs (U1_asRNAs) and tested their efficacy *in vitro*, upon transient co-transfection with RPGR minigene reporter systems in HEK-293T and PC-12 cell lines. We thus identified three chimeric U1_asRNAs that efficiently mediate E9a skipping, correcting the genetic defect. Unexpectedly, the U1-5’antisense construct, which exhibited the highest exon-skipping efficiency in PC-12 cells, induced E9a inclusion in HEK-293T cells, indicating caution in the choice of preclinical model systems when testing RNA splicing-correcting therapies. Our data provide a proof of principle for the application of U1_snRNA exon skipping-based approach to correct splicing defects in RPGR.

## Introduction

In the past several decades, genetic studies have significantly advanced our understanding of inherited retinal dystrophies (IRDs). Retinitis pigmentosa (RP, MIM# 268000) is the most common form of IRD with a prevalence of approximately 1 in 3,000 individuals.^1^ Although clinical symptoms of RP may be similar among patients, the genetic pattern of inheritance is complex. Indeed, RP is a heterogeneous disease associated with sequence variants in more than 71 genes, resulting in different forms of inheritance: autosomal dominant RP (23 genes), autosomal recessive (43 genes) and X-linked RP (XLRP) (5 genes)^2^ (RetNet, see https://sph.uth.edu/RetNet/sum-dis.htm#A-genes). Mutations in the *Retinitis Pigmentosa GTPase Regulator* gene (*RPGR*; MIM# 312610) are responsible for 70% to 80% of XLRP cases.^3–6^ *RPGR* is located on chromosomal region Xp21.1 and spans about 172 kb. RPGR mRNA is found in at least 12 alternatively spliced isoforms and is widely expressed in various tissues (e.g., kidney, brain, retina, lung, and testis), as is its protein. RPGR is essential for the survival of retinal cells, and in humans it is expressed by both rod and cone photoreceptors.^7^ Outside the retina, RPGR has also been detected at the transition zone of primary and motile cilia of airway epithelial cells and centrosomes/basal bodies of cultured cells.^7,8^ Several studies have contributed to defining the ciliary localisation of RPGR and its interacting proteins in the retina.^9^ Some of the alternatively spliced isoforms are retina-specific.^10^ Indeed, the two major subsets of transcripts identified in the retina contain exon ORF15 (RPGR^ORF15^) or exon 19 (RPGR^1–19^).^11,12^ Nevertheless, the mechanism of action of RPGR in photoreceptors is still not well understood. Splicing of RPGR is precisely regulated in a tissue-dependent fashion and mutations in *RPGR* frequently interfere with the expression of alternative transcript isoforms.^13–15^ In the retina, an RPGR splicing isoform has been described to include pseudo-exon 9a (E9a). This exon, 136 bases long, is 418 base pairs (bp) downstream of the 5’ splice site of intron 9. Moreover, a g.26652G>A nucleotide substitution 55 bp upstream of E9a was reported in a patient with a mild RP phenotype.^15^ This substitution affects E9a recognition by the splicing machinery, increasing, by a factor of approximately 3.5, the levels of E9a-containing RPGR transcripts in cone photoreceptors.^15^ The presence of E9a in the mature mRNA results in a premature stop codon and the truncation of the protein. Moreover, E9a RPGR has a peculiar expression pattern, being mainly present in cone photoreceptors rather than in rods. Specifically, this isoform is localised in the inner segment of cones, whereas RPGR is normally expressed in the connective cilium.^15^

Both human proteins RPGR^ORF15^ and RPGR^Ex1-19^ are involved in cilia regulation signaling pathways, but their role is unclear. The RPGR^ORF15^ is a protein with 1152 amino acids and has a Glu-Gly-rich region in the C-terminal domain, while the protein RPGR^Ex1-19^ contains 815 amino acids with an isoprenylation motif at the C-terminus. Both isoforms share exon 1-14 and this suggests that they have some common function. Indeed, RPGR isoforms may compete with each other for the availability of endogenous binding partners. Therefore, maintaining the optimal RPGR^Ex1-19^ / RPGR^ORF15^ ratio has a crucial role in optimal cilia growth and ciliary trafficking regulation.^16^ However, as reported in the literature by Moreno-Leon et al., (2020)^16,17^, both isoforms, RPGR^Ex1-19^ and RPGR^ORF15^ are regulated by independent mechanisms and both of them have different functional properties, different cellular and tissue localization, and different levels of expression during retinal development and maturation.^17^

AAV-based delivery of RPGR^ORF15^ is in advanced clinical development: a clinical trial investigated a gene therapy approach for X-linked RP, via an AAV8 vector-delivered codon-optimized human RPGR^ORF15^ coding sequence (CDS). The initial results of this study were successful and showed an amelioration of the patients’ visual field.^18^ However, as mutations in RPGR may impact several RPGR splicing isoforms at the same time, future gene therapy strategies will need to implement ways to restore all RPGR isoforms, in the correct relative amounts. In this regard, universal gene addition or cell-based therapies, highly promising techniques^19–21^, may not be the optimal choice to treat RPGR mutations. We propose here an exon-skipping strategy, which, besides taking advantage of the endogenous processes guaranteeing a balance among the different RPGR isofroms, would have the additional advantage of being functional only in the cell types in which the target pre-mRNA is expressed.

Intensive research in the field of splicing modulation has led to the development of successful exon-skipping approaches by engineered U1 small nuclear RNA (U1 snRNA) able to mask the mutated sites, thus modulating the aberrant alternative splicing.^22,23^ U1 snRNA recognizes the 5’ splice site and mediates the first step in spliceosome assembly.^24^ Several examples of successful applications of U1 snRNA in therapeutical exon skipping are described in the literature^25–28^. U1 snRNA has been shown to be a useful vector for the stable expression of antisense molecules.^22,28^ The U1 snRNA expression cassettes are small (about 600 bp) and they work efficiently both in *in vitro* and *in vivo* systems and can be delivered as part of lentiviral^29^ or AAV vectors.^25,27,28^

Here, we aimed at developing an exon-skipping U1 antisense snRNA (U1_asRNA) to restore the levels of E9a in RPGR bearing the g.26652G>A deep intronic nucleotide substitution. We designed two single U1_asRNA molecules, U1_3’ and U1_5’, and a double-target U1_3’5’ carrying two distinct antisense sequences targeted against two splicing-regulating regions. This latter strategy has already been used to increase exon skipping efficiency.^25–27^ We tested whether the three chimeric constructs were able to modulate E9a skipping in HEK-293T and PC-12 cell lines, by co-transfecting them with a RPGR minigene reporter system, recapitulating mRNA RPGR expression observed in patients (MINI mut) and unaffected control individuals (MINI wt).

Our results demonstrate that a new therapeutic strategy based on U1_snRNA molecules can efficiently be used to correct the splicing of *RPGR* transcripts. Additionally, the finding that U1_5’asRNA induces exon E9a skipping in PC-12 cells and inclusion in the HEK-293T cells, calls for caution in the choice of the preclinical model systems used to test U1_asRNAs.

## Results

### Computational analysis of splicing-relevant sequences in RPGR exon 9a

In order to understand how to modulate splicing of E9a, we undertook a bioinformatics analysis of sequences essential for the recognition of the alternative exon. We also investigated the effect of the g.26652G>A nucleotide substitution on these sequences.

The first analysis carried out by using NNSPLICE 0.9 program^30^ (https://omictools.com/nnsplice-tool), showed that E9a has a weak consensus sequence at its 3’ splice site, indicating that a Splicing Enhancer might assist E9a splicing, whereas its 5’ splice site is similar to the consensus sequence (**Table 1**). Additionally, we analysed whether the nucleotide substitution might create a new branch site with a stronger consensus. However, using the branch point analysis bioinformatic program “Human Splicing Finder”^31^ (http://www.umd.be/HSF/), we found no consensus change in the branch sites pattern of intron 9 (data not shown).

**Table 1.**
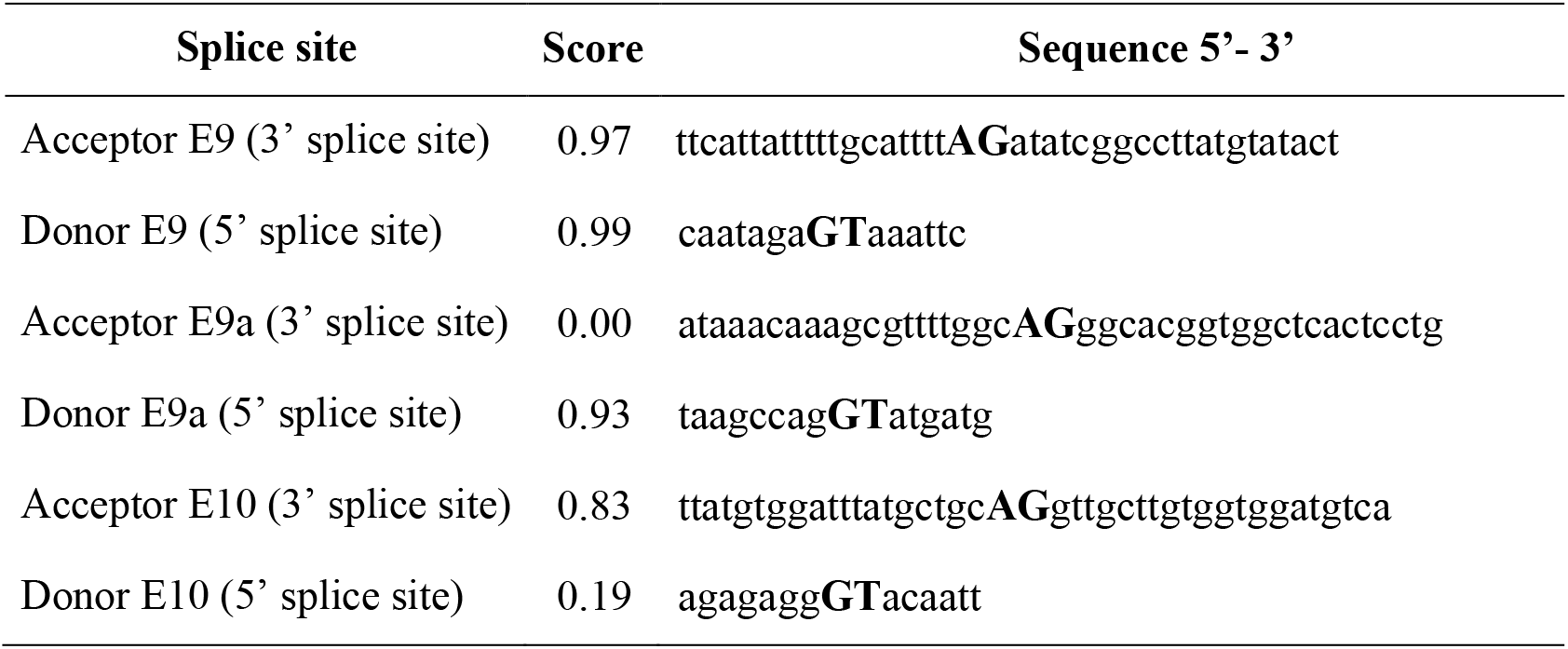
Splice site scores of RPGR sequence from exon 9 to 10. According to NNSPLICE 0.9, higher scores indicate stronger splice sites. A perfect match to the splice site consensus sequence has score 1.

Identification of a defined Exon Splicing Enhancer (ESE)^32^ sequence on E9a would provide an optional target site for the design of chimeric U1_snRNAs able to mediate E9a skipping. SpliceAid site predictions^33^ (http://www.introni.it/splicing.html) for E9a did not show the presence of a clear and defined ESE along its sequence, since both splice-enhancing and splice-silencing RNPs are predicted to bind to the same regions (**Supplemental Figure S1A**). Moreover, a comparative analysis of mutant and wild-type intronic region sequences (100 nt upstream of E9a) did not highlight any possible involvement of intronic splice enhancers or silencers in E9a splicing (data not shown). Nonetheless, a secondary structure prediction (**Supplemental Figure S1B**) indicates that the g26652G>A intronic nucleotide substitution destabilizes a 13-bp-long stem-loop in the intron, while not affecting the accessibility of the E9a acceptor site.

Our bioinformatic analyses suggested that E9a does not have any evident ESEs. Moreover, the nucleotide substitution site does not seem to represent a valid target for inducing exon skipping. Accordingly, we decided to target E9a splice sites. Indeed, masking these sites should result in E9a skipping as a consequence of the splicing machinery failure to recognize these critical *cis*-acting sequences.

### Analysis of exon 9a inclusion in RPGR mRNA in cells in culture

We aimed at analyzing exon 9a inclusion in RPGR mRNA in two independent cell models, to add robustness to the analysis. Moreover, the chosen cell lines had to be easy to grow, to maintain and to transfect. Finally, the cell models should recapitulate as much as possible the repertoire of auxiliary splicing factors typical of photoreceptors and other neurons. Alternative splicing is indeed a mechanism increasing gene-expression diversity which is widely used by neurons, and exon-skipping therapeutical strategies should be tested taking the tissue-specific peculiarities of this mechanism into account.^23^

High levels of expression of both *RPGR* mRNA and protein have been reported in the human adrenal gland medulla (https://www.proteinatlas.org/ENSG00000156313-RPGR) that has a neural crest origin as do other neuronal types. We, therefore, chose to use two different adrenomedullary cell lines: HEK-293T and PC-12.

HEK-293T (fetal human embryonic kidney cells)^34^ derive from HEK-293 that, despite their name, have presumably been established from a human (female) embryonic adrenal precursor cell.^35^ HEK-293T cells are hypotriploid and carry three copies of the X chromosome.

PC-12 cells derive from a pheochromocytoma in a (male) rat adrenal medulla^36^ and grow in culture as undifferentiated neuroblasts. They are a commonly employed model system for studies of neuronal development and function and are relatively easy to passage, culture and transfect.^37^

The use of PC-12 cells of rat origin, additionally, allows to enquire whether the therapeutical strategy could be tested in rodent models of the disease, in the future.

By semiquantitative RT-PCR performed with primers annealing to exons 7 and 10 of RPGR mRNA, we evaluated the level of endogenous RPGR expression in HEK-293T and PC-12. We showed that, in both cell lines, endogenous *RPGR* mRNA was detectable in the form of E9a-, while the E9a+ isoform was not present (**Figure 1**). RPGR mRNA was more abundant in HEK-293T than in PC-12 cells.

**Figure 1.**
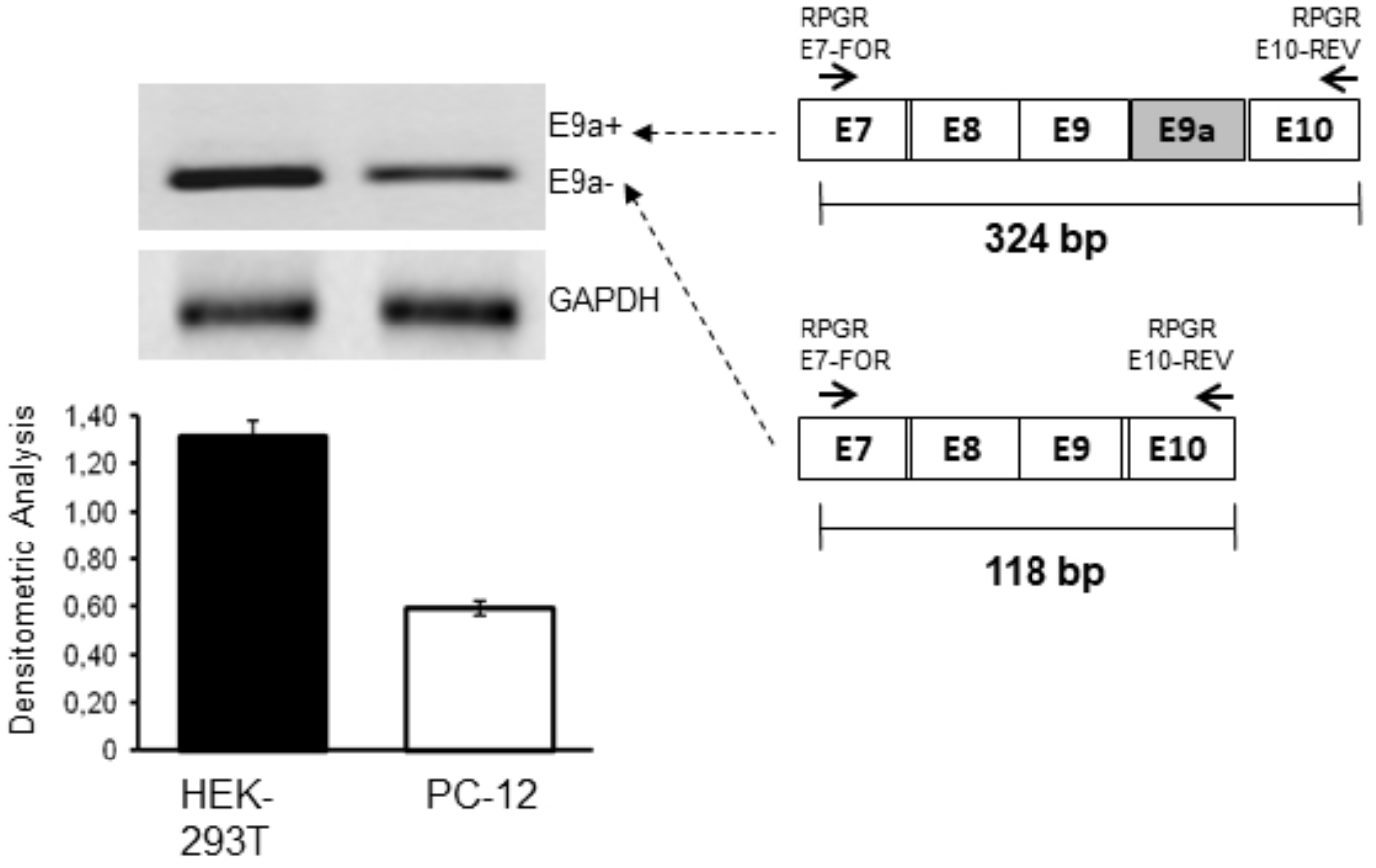
Analysis of exon 9a inclusion in RPGR mRNA in HEK-293T and PC-12 cells in culture. Semiquantitative RT-PCR analysis was performed with primers annealing to exon 7 (forward) and exon 10 (reverse) of human and rat RPGR mRNA, respectively. Lane 1: HEK-293T cells, Lane 2: PC-12 cells. As depicted in the schematic diagram to the right, the presence of E9a would yield a 324-bp-long amplicon (E9a+), while the amplification of RPGR transcript devoid of E9a produces an E9a-118-bp-long amplicon. GAPDH was used as a housekeeping gene. One representative gel of three is shown. Only E9a-bands were detected. Their intensities were measured by densitometric analysis and reported in the histogram. Values are represented as means ± S.D (n=3).

### Reporter minigenes recapitulate RPGR exon 9a alternative splicing

To study E9a skipping, *in vitro*, we constructed reporter systems by cloning the RPGR genomic region containing exon 9, the entire intron 9 and exon 10 in the expression vector pCDNA3. We produced both the RPGR wild-type reporter (MINI wt) and the mutant one (MINI mut) bearing the g.26652G>A mutation located 55 bp upstream of the E9a 3’splice site (**Figure 2A**). We expect the RPGR mutant reporter to generate higher levels of transcripts containing exon 9a compared to the wt reporter, as observed in RP patients.^15^

**Figure 2.**
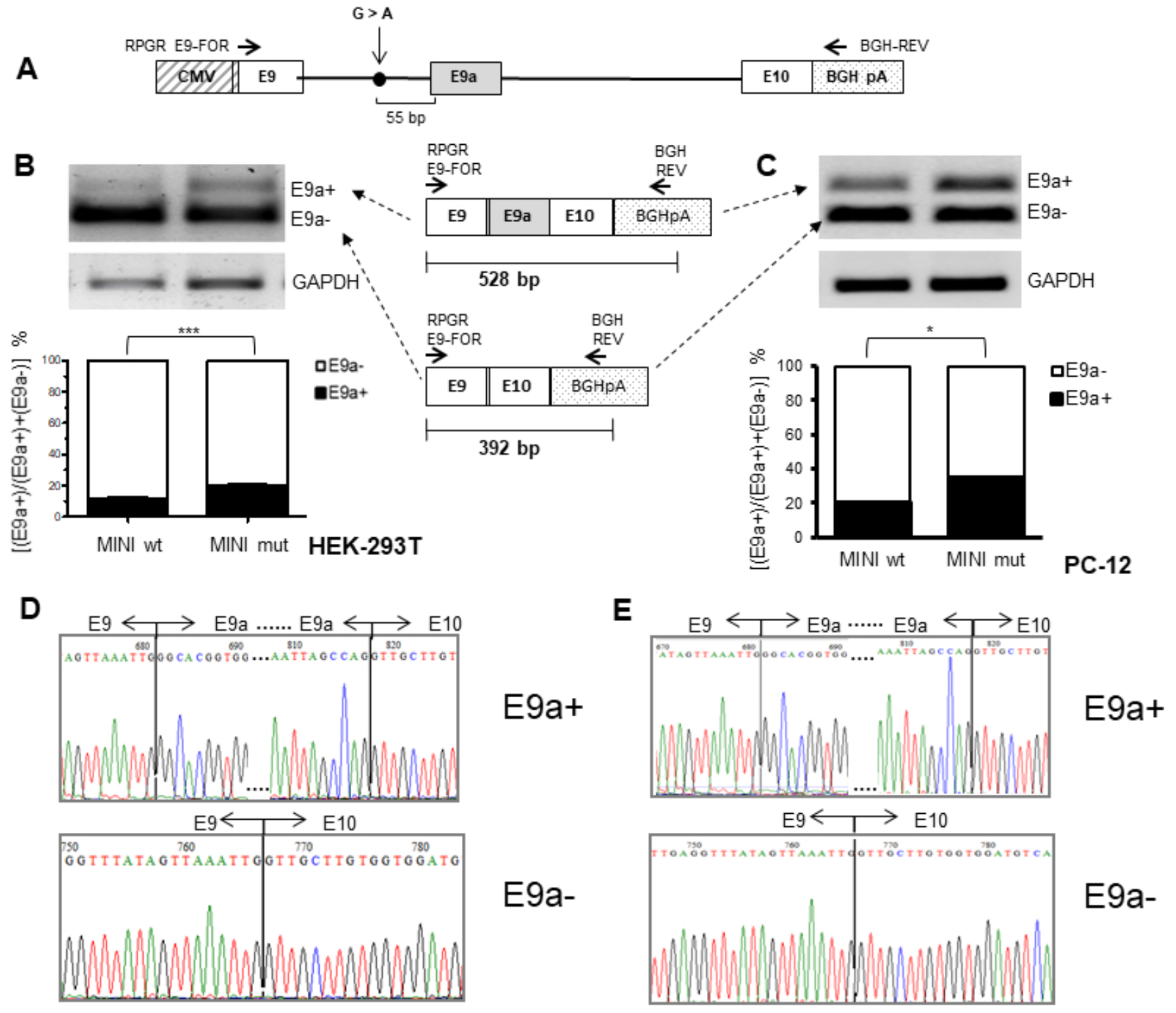
Setting up a minigene reporter system of RPGR E9-E10 splicing and E9a inclusion-inducing mutations. **(a)** Schematic representation of the RPGR minigene construct (not in scale). The position of the mutation is indicated by a dot, while horizontal arrows represent primers for RT-PCR analysis. The endogenous RPGR mRNA is not amplified by this primer pair, as the reverse primer anneals to the reporter-specific portion of the RNA transcribed from the BGH polyA cassette. The alternative splicing of exon 9a leads to the production of two different splice isoforms (E9a+ and E9a-). As depicted in the schematic diagram below, the presence of E9a in the reporter RPGR transcript yields a 528-bp-long amplicon (E9a+), while the amplification of the reporter RPGR transcript devoid of E9a produces a 392-bp-long amplicon (E9a-). **(b)** RT-PCR analysis of RPGR mRNA levels on RNA from HEK-293T cells transfected with wild-type minigene (MINI wt, lane 1) and mutant minigene (MINI mut, lane 2). The analysis has been performed in triplicate, and one representative gel is shown. The histogram represents the densitometric analysis of the bands relative to the two different isoforms (E9a+ and E9a-) normalized to GAPDH (p<0.001). Data are shown as mean ± S.D (n=3). **(c)** RT-PCR on RNA from transfected PC-12 cells. Lane 1: wild-type minigene (MINI wt), Lane 2: mutant minigene (MINI mut). The histogram represents the densitometric analysis of the bands relative to the two different isoforms (E9a+ and E9a-) normalized to GAPDH (p<0.05). Data are shown as mean ± S.D (n=3). **(d) and (e)** E9a+ and E9a-bands from lanes 2, of the gels in panels (b) and (c), respectively were eluted and sequenced. Chromatograms show the correct fusion between exons 9 and 9a, and exons 9a and 10, in the E9a+ band, and exons 9 and 10 in the E9a-band.

To test if the reporter constructs were able to recapitulate alternative splicing of E9a in presence or absence of the g.26652G>A nucleotide substitution, we transiently transfected either MINI wt or MINI mut in HEK-293T and PC-12 cells. By semiquantitative RT-PCR, using primers targeting RPGR exon 9 and the BGHpA region (**Figure 2A**), we observed that in both cell lines the nucleotide substitution was able to increase E9a levels of about two-fold compared to the wild-type (**Figures 2B and 2C**). PCR products were analysed to confirm the exact sequence of the two mRNA RPGR isoforms, with and without E9a, in HEK-293T and PC-12 cells transfected with MINI mut reporters (**Figure 2D and 2E**, respectively).

### Design of antisense U1_snRNAs to induce exon skipping of RPGR exon9a

To induce skipping of E9a in the context of our RPGR minigene reporters, we designed and generated three chimeric constructs in the backbone of the U1_snRNA.

The nucleotide sequence required for the recognition of the 5’ splice site (positions 3 - 10 at the 5’-end of U1 snRNA) was substituted with an antisense sequence complementary either to the 3’ splice site (U1_3’), 5’ splice site (U1_5’) or both splice sites simultaneously (U1_3’5’) (**Figure 3A**). This latter construct was created because, according to literature, there is a higher efficiency of exon skipping by employing double-target U1 asRNAs.^25–27^ As a negative control, we additionally cloned an U1_Scramble construct, bearing a 23-nt-long sequence (**Figure 3A**) with no targets in mammalian cells.

**Figure 3.**
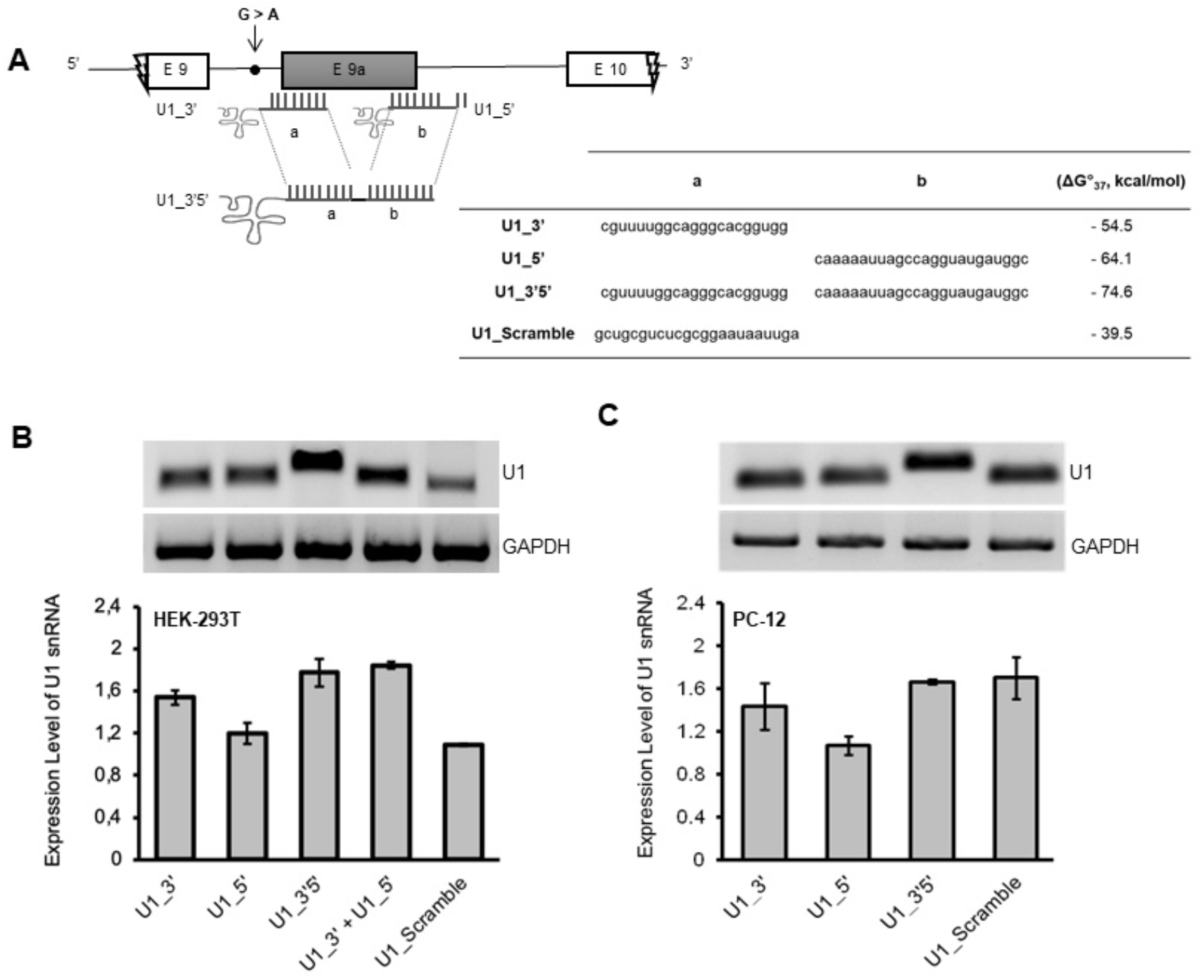
Validation of chimeric antisense U1 snRNA constructs expression. **(a)** Schematic representation of chimeric U1 snRNAs with splicing correction activity on the minigene model of RPGR (not in scale). The table summarizes the target regions of the different constructs, the scramble sequence (5’-to-3’ direction) and the values of the hybridization energy of Chimeric U1_snRNAs/RPGR pre-mRNA (ΔG°_37_, Kcal/mol), predicted by using DuplexFold tool. **(b and c)** Expression of chimeric antisense U1 constructs in HEK-293T (b) and PC-12 (c) was assayed by RT-PCR, using primers U1+130 Rev and the appropriate U1-RPGRFor (see materials and methods). One representative gel of three is shown in both (b) and (c). The histogram shows the densitometric analysis as mean ± S.D (n=3).

Since the different antisense constructs, U1_3’, U1_5’, U1_3’5’ and U1_Scramble, considerably extended the length of the U1_snRNA, we checked both integrity and expression levels of U1 antisense RNAs after co-transfection in HEK-293T and PC-12 cells. By performing semiquantitative RT-PCR as described in materials and methods, we observed the presence of specific amplification products, confirming that all designed molecules were present and expressed at comparable levels (**Figures 3B and 3C**). As expected, the sizes of the amplicons (around 100 bp) slightly varied, depending on the antisense sequence introduced and on the different forward primer used.

### Antisense U1_snRNAs restore RPGR transcript expression patterns

In order to assess the feasibility of E9a skipping, we carried out co-transfection experiments by using the U1_3’, U1_5’, U1_3’5’, U1_3’ + U1_5’ and U1_Scramble constructs in combination with the RPGR MINI mut minigenes, in HEK293 cells.

When using MINI mut, we observed a reduction of E9a (E9a+) mRNAs of about 50% using U1_3’ or U1_3’5’ in HEK-293T cells, when compared to the transfection of the minigene alone. Surprisingly, the U1_5’ led to a 130% increase in E9a+ transcripts (**Figure 4A**).

**Figure 4.**
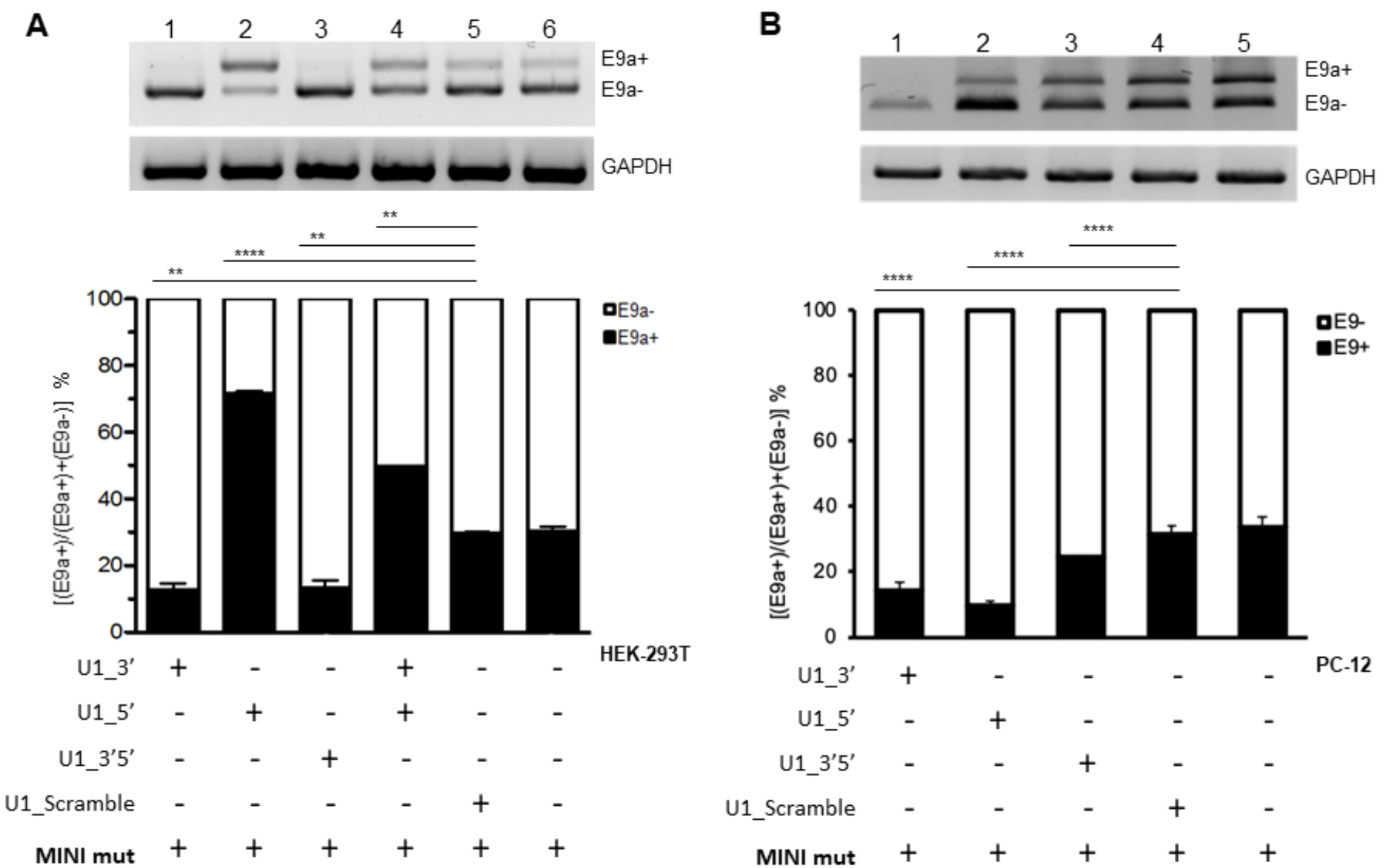
Chimeric antisense U1 snRNA-induced exon 9a skipping in HEK-293T and PC-12 cell lines transfected with RPGR mutant minigene. **(a)** Semiquantitative RT-PCR of RNA from HEK-293T cells transfected with RPGR mutant minigene (MIN mut) alone or in combination with chimeric U1_snRNAs plasmids. E9a reduction using U1_3’ or U1_3’5’ is significant (p<0.01) compared to U1_Scramble. **(b)** Semiquantitative RT-PCR of RNA from PC-12 cells transfected with RPGR mutant minigene (MINI mut) alone or in combination with chimeric U1 snRNAs. E9a reduction using any of the three U1 constructs is significant (p<0.001) compared to U1_Scramble. One representative gel of three is shown in both (a) and (b). Densitometric analysis of E9a+ and E9a-amplicons, from three independent experiments, is shown for both cell lines. GAPDH is used as an internal control. Data are shown as mean ± S.D (n=3). Statistical analysis was performed with a global statistical test and is reported in Supplemental Table S1 (p-value: **P < 0.01; ***P < 0.001).

Additionally, when both constructs U1_3’ and U1_5’ were cotransfected, E9a (E9a+) inclusion increased in mature mRNA, suggesting that U1_5’ has a dominant effect on U1_3’. The fact that the total amount of U1_asRNA was maintained (transfecting only half of the amount of U1_5’ compared to when it was transfected alone) explains why exon inclusion was 50% lower than that obtained by transfecting U1_5’ alone.

The U1_Scramble control did not cause any significant alteration in the RPGR expression, as expected. Overall, our results indicate that the U1_3’ and U1_3’5’ constructs (**Figure 4A**) induce the exon skipping correction for g.26652G>A nucleotide substitution.

Statistical analysis of these data was performed by a global statistical test (two-way ANOVA) and detailed data are reported in **Supplemental Table S1**.

All the E9a+ and E9a-bands in Figure 4A were eluted and sequenced, confirming the correct joining of exons 9 and 10, or exons 9, 9a and 10, respectively. In particular, sequencing of bands in lanes 2 and 4 confirmed that the lower band consists in the transcript devoid of exon 9a, and the upper band results from the correct joining of exon 9a in between exon 9 and exon 10 (**Supplemental Figure S2**).

We then selected one of our best-performing chimeric U1 snRNA, U1_3’, and tested it at decreasing concentrations (200, 66, 20 ng) over RPGR MINI mut transfected HEK-293T cells, to assess if its effect was dose-dependent. Semiquantitative RT-PCR assay clearly showed that the U1_Scramble does not affect exon 9a levels at the different tested doses, on the contrary U1_3’ also showed measurable E9a skipping at lower doses (**Supplemental Figure S3**).

To test the exon skipping effects in a different cellular environment, we analysed total RNA isolated from PC-12 cells transfected with the RPGR MINI mut alone or in combination with our U1_snRNAs (**Figure 4B**). All three designed U1_asRNAs (U1_3’, U1_5’and U1_3’5’) induced E9a skipping, albeit at different levels. Indeed, the constructs’ ability to reduce the inclusion of the E9a depended on the targeted sequence and ranged from ~70% reduction for U1_5’ to ~20% reduction for U1_3’5’ compared to the MINI mut alone. The U1_Scramble did not influence RPGR E9a splicing, as expected (**Figure 4B**).

Sequence analysis of both E9a+ and E9a-PCR isoform products was performed on all bands shown in Figure 4B and showed restoration of the reading frames after treatment with U1_snRNA constructs and the presence of the exact junction between exon 9 and exon 10 in the products correlated to skipping. Supplemental Figure S4 reports the chromatograms relative to lane 2 of **Figure 4B**.

Thus, interestingly, we observed that the efficiency of our constructs is not only dependent on the target sequence but also on the cell type. In contrast to the results obtained in HEK-293T cells (**Figure 4A**), U1_5’ led to efficient E9a skipping in PC-12 cells (**Figure 4B**).

The U1_asRNAs also recognize the sequence of wild-type RPGR pre-mRNA. However, semiquantitative RT-PCR analysis carried out using the RPGR MINI wt with the U1_snRNA constructs (U1_3’, U1_5’and U1_3’5’) did not show any significant variation on endogenous RPGR mRNA expression levels for both HEK-293T and PC-12 cell lines, compared with the non-treated transfected RPGR wild-type (p=n.s.; n=3) (**Supplemental Figures S5A and S5B**, respectively).

### Chimeric antisense U1 asRNAs are able to form snRNPs complexes

To test whether the U1_asRNAs constructs are able to assemble in functional U1 snRNPs, upon co-transfection of HEK-293T cells with the RPGR MINI mut and U1_3’, U1_5’ and U1_Scramble constructs, we performed an RNA ImmunoPrecipitation (RIP) assay.^38^ Nuclear extracts were prepared from transfected cells (**Figure 5A**) and immunoprecipitated with an anti U1-70 K antibody. This antibody recognizes the U1 snRNP 70K protein that, by interacting and binding to the stem-loop I of U1 snRNA in the first step of the spliceosome assembly, mediates the pre-mRNA splicing behaving as a *trans*-acting factor (**Figure 5B**).^27^ Immunoprecipitated RNP complexes were then dissociated into RNA and protein. The RNA was isolated and checked for integrity and length by semiquantitative RT-PCR. As expected, PCR products were detected in Nuclear Extracts (NE) and enriched in the samples immunoprecipitated with anti-U1-70K (IPP) (**Figure 5C**). PCR products of U1_snRNAs were barely detectable in the sample treated with normal rabbit IgG conjugated to beads, (IgG), Input and no template control (**Figure 5C**). These results indicate that both U1_3’ and U1_5’ antisense U1_snRNAs constructs are able to assemble into a functional U1 snRNP.

**Figure 5.**
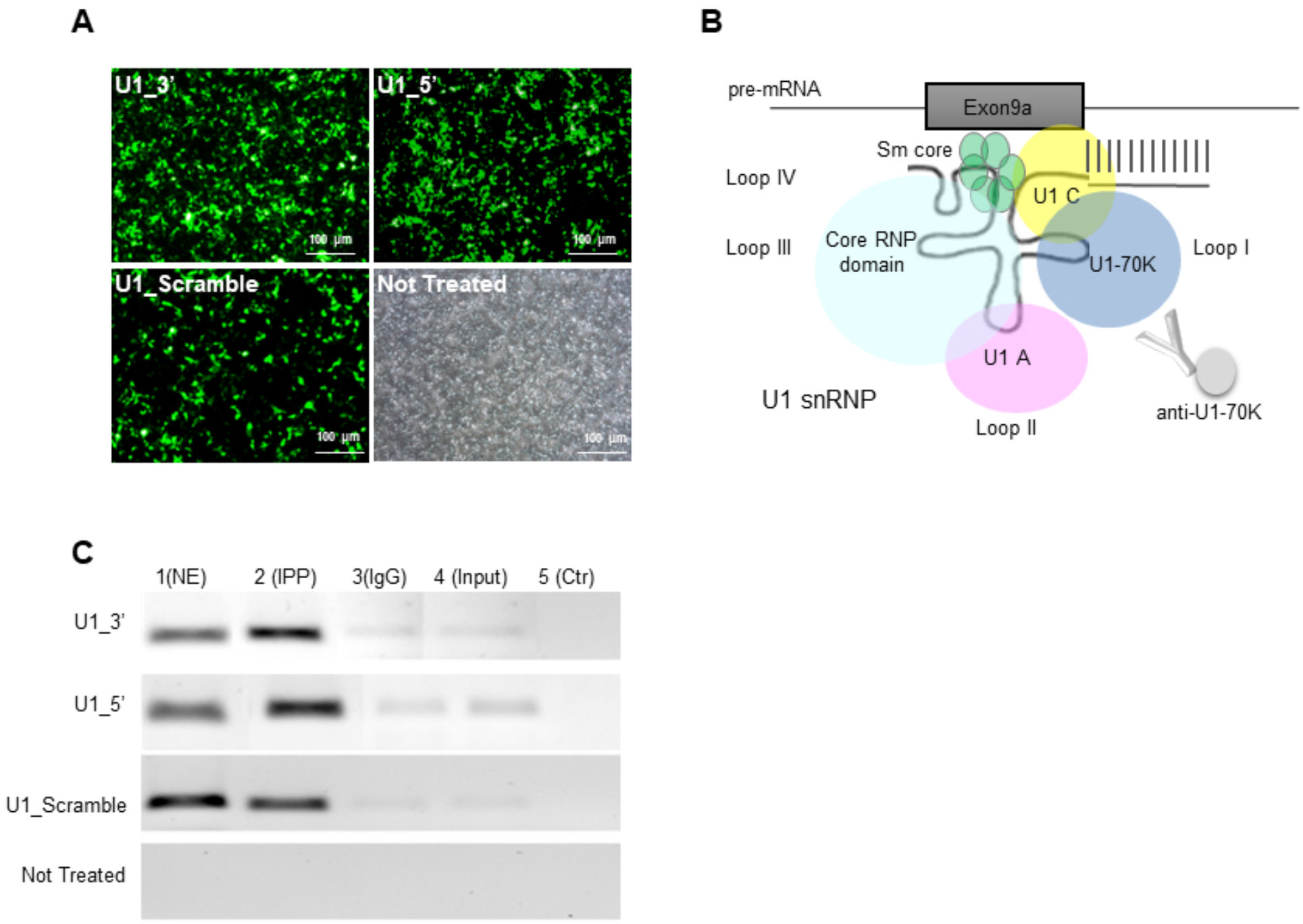
Chimeric U1_5’ and U1_3’ assemble with U1-70k ribonucleoprotein in transfected HEK-293T cells. **(a)** Representative images of HEK-293T cells 48 hours after co-transfection with the RPGR mutant minigene and the different chimeric U1_asRNAs constructs (Zeiss Axio Observer Z1 microscope; Scale bar: 100μm). **(b)** Schematic representation of the U1 snRNP particle (not in scale). **(c)** Semiquantitative RT-PCR of the immunoprecipitated RNA using anti U1-70k antibodies was performed with a U1+130 primer and specific U1 primers, as described in materials and methods. **NE**: Nuclear extracts; **IPP**: NE sample + U1-70K antibody; **IgG**: NE sample + IgGs; **Input**: NE sample + Beads; **Ctr**: No Template PCR control.

## Discussion

Most eukaryotic genes contain pseudoexons, sequences that look like perfect exons but are ignored by the splicing machinery based on rules that are yet to be fully understood. In recent years, research has highlighted roles for pseudoexons not only as regulators of the splicing process but also as possible causes of human diseases.^39^

Here, we focused on an intronic mutation that affects pseudoexon splicing but does not alter the consensus sequence of the pseudoexon splice sites. This type of mutation acts mainly by deleting or creating Intronic Splicing Silencers (ISS) or Enhancers (ISE) or changing the strength of the consensus sequence of the branch point.^40–42^ There are two advantages in targeting splicing mutations instead of mutations leading to missense or frameshift. Firstly, since these mutations influence splicing, the achievement of correction of the altered splicing pattern is sufficient to obtain a therapeutic benefit. Indeed, if constitutive exons won’t be skipped, functional analysis of the rescued proteins is not needed. Secondly, if the mutations affect splicing by creating new *cis*-acting sequences, the disease can be quickly addressed by identifying and masking those sequences with antisense oligonucleotides approaches. For these reasons, we selected the intronic nucleotide substitution g.26652G>A of the RPGR gene^15^ as an optimal target for applying splicing-correction approaches in the retina.

Neidhardt and colleagues^15^ reported that the nucleotide substitution g.26652G>A in the RPGR gene increases the levels of the alternatively spliced exon 9a non-sense pseudoexon. Our bioinformatic analyses were not conclusive about whether this nucleotide substitution causes an increase of the alternatively spliced E9a RPGR mRNA through the generation or removal of *cis*-acting sequences.

Nevertheless, we designed chimeric U1_snRNAs, (U1_5’, U1_3’, U1_3’5’ and U1_Scramble), directed towards either splice sites of E9a or both splice sites at the same time (double U1_asRNA), aiming at reducing the splice site recognition in the presence of the G>A nucleotide alteration. U1_snRNAs typically recognise the 5’ splice site by the binding of their 5’ end to the - 3/+ 6 exon/intron consensus sequence. This binding needs to be reversible since U1 snRNA has to detach from the 5’ splice site so that the splicing reaction can proceed further. To increase the specificity and affinity of U1 to its target site, we increased the length of the region recognized by our chimeric U1_asRNAs to 21, 24 or 45 nucleotides.

Using RNAfold, it was possible to predict the secondary structure and the ΔG°_37_ free energy of both wild-type and mutant RPGR pre-mRNA sequences. Therefore, the Chimeric U1_snRNAs were designed to take into consideration the folding energy (data not shown) and the binding energy between the U1_snRNA Chimeras and RPGR pre-mRNA target regions (Figure 3A). Comparing the average of binding Gibbs free energy (ΔG°_37_, kcal/mol) of endogenous U1_snRNA with our chimeric U1_snRNAs, we observed that the ΔG°_37_, (kcal/mol) values of our U1_snRNA were lower than the endogenous U1_snRNA (Figure 3A). Therefore, the computational analysis confirmed the efficient binding of our chimeric U1_snRNAs to the RPGR pre-mRNA target sequence, suggesting that they could play a role, *in vitro*, in E9a exon-skipping.

To recapitulate the wild-type and mutant conditions *in vitro*, we generated minigene reporter systems.^43–45^ We designed a wild-type minigene (MINI wt), and one carrying the g.26652G>A mutation (MINI mut) (**Figure 2A**). These constructs were transfected either in Human Embryonic Kidney (HEK-293T) cells or rat pheochromocytoma (PC-12) cells. Both cells express the endogenous RPGR mRNA exclusively without the E9a pseudogene (**Figure 1**). Increased levels of E9a transcripts were detected upon MINI mut transfection compared to cells in which the MINI wt was transfected, in both HEK-293T and PC-12 cells, recapitulating the splicing pattern of patients’ cells affected by the g.26652G>A mutation.^15^

Neidhardt and colleagues^15^ have estimated that in human retinas from healthy individuals approximately 4% of RPGR transcripts bear exon 9a (Figure 1C in Neidhardt et al., 2007).^15^

In our hands, upon transfection of our wild-type RPGR minigene, exon 9a is included in 12% of RPGR minigene-derived transcripts in HEK-293T and in 18% of said transcripts in PC12 cells (**Figure 2B and 2C**, our manuscript).

These observations are in line with the general understanding that alternative splicing is not a black-and-white process and small amounts of non-sense alternative (pseudo-)exons are often spliced in gene transcripts, possibly as part of a finely tuned post-transcriptional regulation of gene expression.^15^

Neidhardt and colleagues (2007) describe a mutation in the RPGR gene (g.26652 G>A) which correlates with X-linked Retinitis Pigmentosa and significantly increases exon 9a inclusion in RPGR transcripts. When comparing RPGR transcripts in blood cells from the affected individual to those of his two unaffected brothers, Neidhardt and co-workers report a 3-times higher level of exon 9a-containing RPGR transcript.

Therefore, our mutant minigene system recapitulates the molecular defect observed in the patient (i.e. the increase of exon9a-containing transcripts). In fact, upon transfection of an RPGR minigene bearing mutation g.26652 G>A (“miniMUT”), exon 9a is included in 20% of RPGR minigene-derived transcripts in HEK-293T (Fig. 2B) and in 30% of said transcripts in PC12 cells (**Figure 2C**).

By testing U1_asRNAs against the RPGR minigene system, we observed that a chimeric U1_asRNA directed towards the 3’ splice site (U1_3’) in HEK-293T and PC-12 cells was able to achieve a significant E9a exon skipping (**Figure 4**).

Interestingly, we observed a different effect of the U1_5’ construct activity. We designed this construct with a 24 bp long antisense sequence in order to have a stable hetero-duplex conformation and to promote masking of the splice site, in line with results reported in the literature.^26,47^

The U1_snRNA directed towards the 5’ splice site unexpectedly caused a significant increase in E9a levels when tested in the HEK-293T (**Figure 4A**). Indeed, sequence analysis performed on the E9a+ PCR product (**Supplemental Figure S2**) confirmed that the fragment corresponded to the sequence containing E9, E9a, and E10 perfectly joined.

On the contrary, the U1_5’ was as capable as the U1_3’ of skipping exon E9a in PC-12 cells, restoring the normal expression of the alternatively spliced RPGR mRNA (**Figure 4B**). These findings bring into question why this approach does not work in all cell types in the same way. Additionally, it has to be noted that U1_5’ does not induce E9a inclusion in MINI wt transfected HEK-293T cells (**Supplemental Figure S5A**).

Overall, it seems that g. 26652G>A mutation facilitates E9a inclusion and the U1_5’ in HEK-293T cells acts over the mutated minigene similarly to adapted U1 snRNAs, which are typically designed with shorter complementarity (around ten base pairs).^48–50^ It was demonstrated, indeed, that when complementarity between U1_snRNA and 5’splice site is extended to 11 nt, 5’ splice site recognition is increased.^51^ Notably, RNA immunoprecipitation (RIP) experiments were performed to check the capability of our U1_asRNA chimeras to bind the U1-70K protein, proving that they can form stable and functional snRNPs (**Figure 5**).

We also analyzed the effect of the double antisense construct U1_3’5’ on both HEK-293T and PC-12 cell lines. This construct led to splicing correction in HEK-293T at a comparable level of that induced by U1_3’, reducing the percentage of E9a-containing transcripts to half (**Figure 4A**) while in PC-12 U1_3’5’ was less efficient than U1_3’, just reducing the percentage of E9a-containing transcripts to ¾ relative to U1_Scramble (**Figure 4B**).

Interestingly, while evaluating the combined effect of U1_3’ and U1_5’ antisense constructs, we observed an increase of E9a levels in between those observed with each of the two chimeric RNA (**Figure 4A**). This result suggests that U1_5’ exon-inclusion effect overtakes U1_3’ exons-skipping effect. The observed levels of E9a inclusion were 50% lower than that obtained by transfecting U1_5’ alone, in line with the reduced amounts of U1_5’construct transfected.

Our study provides evidence that both our chimeric constructs U1_3’ and U1_3’5’, tested in HEK-293T and PC-12 cell lines, are efficient at inducing exon skipping of RPGR Exon 9 in the presence of nucleotide substitution g.26652G> A. However, one of our constructs, U1_5’, induces exon skipping or exon inclusion, depending on the cellular background.

In speculating why U1_5’ has opposite effects in HEK293T versus PC-12 cells we have reasoned that the same mutated minigene is spliced to include E9a with higher efficiency in PC12 cells than HEK293T cells, as evident in Figure 2B and 2C. The reason of this higher efficiency might lie in the better binding of E9a splice sites by U1snRNP or accessory proteins, or their higher levels. In line with this reasoning, the introduction of U1_5’ asRNA in PC-12 cells interferes with this E9a splicing mechanism by inhibiting U1snRNP binding by virtue of its extended complementarity to the donor site. On the other hand, in HEK-293T cells, U1snRNP is possibly less efficient or less abundant, and the exogenous U1_5’ asRNA might replace the endogenous U1 snRNA, therefore acting as an adapted U1, attracting the U1 snRNP components to the donor site and ultimately stimulating E9a exon inclusion.

Taken together, these results suggest that a new therapeutic strategy based on U1_snRNA molecules could efficiently be used to restore the physiological RPGR gene splicing, but call for caution in the choice of the model system used to study its efficiency.

The results presented here constitute proof-of-concept data that should be followed by further preclinical trials in patient-derived cells and in animal models, to test efficacy in correcting the genetic defect, with more emphasis on the correction of the genotype-phenotype pattern. However unfortunately, to our best knowledge, no patient-derived cell model of RPGR Exon 9a inclusion is available at the moment that could be used for further studies, nor any animal model.

The U1 promoter/U1_asRNA cassette is relatively small (ca. 600 bp) and can easily be accommodated in the limited capacity of AAV vectors. Based on our previous experience in a mouse model of Duchenne Muscular Dystrophy, exon-skipping U1asRNAs vectored by AAV show a good distribution and are effective and stable.^25,28^ Upon a successful preclinical proof-of-concept study, the described therapeutic approach could be translated into clinical trials via subretinal injection^52,53^ that is performed by administrating the therapeutic molecule between the photoreceptor cell layer and the retinal pigment epithelium (RPE).

## Materials and methods

### Computational analysis of RPGR exon 9a splicing

Splice sites were explored by analyzing the RPGR genomic sequences from exon 9 to 10 with the NNSPLICE 0.9 program^30^ (https://omictools.com/nnsplice-tool). The resulting scores for the different splice sites were reported (**Table 1**). Moreover, the branch point analysis tool was conducted on intron 9 using the Human Splicing Finder (http://www.umd.be/HSF/).^31^

### Construction of RPGR Reporter Minigenes

To generate minigene constructs, we cloned genomic sequences spanning a region from intron 8 to 10 of the *RPGR* gene into the mammalian expression vector pcDNA3 (Life Technologies, Carlsbad, CA, USA). RPGR wild-type minigene construct was generated (**Figure 2A**) by amplification from a pool of human female genomic DNAs (Promega, Milan, Italy) using the following primers:

RPGR-For: 5’-ctaggtacccacagagaccatagagagtg-3’ and
RPGR-Rev: 5’ -ctactcgagaagtttgttagcactcaactctaa-3’

PCR amplification was performed in a final volume of 50 μl, using the Cloned Pfu DNA polymerase (Agilent Technology, Santa Clara, CA, USA), according to the manufacturer’s protocol.

The PCR products and the pcDNA3 vector (Life Technologies, Carlsbad, CA, USA) were cut with endonucleases KpnI and XhoI (New England Biolabs, MA, USA) and purified using the QiAquick Gel Extraction Kit according to the manufacturer’s protocol (Qiagen, Hilden, Germany). The digested fragments were cloned into the pcDNA3 vector using T4 DNA ligase (New England Biolabs, MA, USA), according to the manufacturer’s protocol.

The generated RPGR minigene wild-type (MINI wt) was used as the template for mutant RPGR minigene production (MINI mut). To introduce the g.26652G>A mutation we used the Quick-change II XL Site-directed Mutagenesis Kit (Agilent Technologies, USA), according to the manufacturer’s protocol. The PCR amplification was performed in a final volume of 50 μl in a reaction mixture containing the RPGR wild-type minigene, as template, and the following specific primers: mutRPGR-For: 5’-gctgaattaaatgttaaactctcaaatcctgcacaacag-3’ and mutRPGR-Rev: 5’-ctgttgtgcaggatttgagagtttaacatttaattcagc-3’. The sequence of both wild-type and mutant minigenes and the exact point mutation position into the RPGR mutant minigene were verified by sequencing (BMR Genomics, Padova).

### Cloning of U1 expression constructs

Four chimeric U1_snRNA constructs were obtained by ligation of two different inverse polymerase chain reaction (PCR) products. The first containing the U1 promoter, and the second containing the U1 sequence plus the chimeric sequence, as described in Denti et al. (2006).^54^ To generate the U1_5’, U1_3’ and U1_Scramble constructs, a first PCR was performed on the human U1 snRNA gene using the forward primer U1cas-up For: 5’-ctagctagcggtaaggaccagcttctttg-3’ and three different reverse primers: U1_5’RPGR Rev: 5’-caaaaattagccaggtatgatggcatgagatcttgggcctctgc-3’,

U1_3’RPGR Rev: 5’-cgttttggcagggcacggtggatgagatcttgggcctctgc-3’, or U1_Scramble RPGR Rev: 5’-tcaattattccgcgagacgcagcatgagatcttgggcctctgc-3’.

All PCR assays were carried out in 50 μl final volume in a reaction mixture containing 1X of Cloned *Pfu* DNA polymerase Taq (Agilent Technology, Santa Clara, CA, USA), according to the manufacturer’s protocol. Each of the three PCR fragments produced was then ligated with a PCR product from the human U1 snRNA gene using the U1-univ For primer: 5’-ggcaggggagataccatgatc-3’ and U1Cas-down Rev primer: 5’-ctagctagcggttagcgtacagtctac-3’.

To generate the U1_5’3’ constructs, a PCR product, obtained using primers U1Cas-up For and U1 5’RPGR Rev, was ligated with another PCR product, generated using primers U1 3’RPGR-II For: 5’-ccaccgtgccctgccaaaacgggcaggggagataccatgatc-3’ and U1Cas-down Rev. Ligation was performed in a final volume of 20 μl by incubation at 16°C for 3 hours, according to the manufacturer’s protocol (T4 DNA ligase, New England BioLabs, NEB, Ipswich, MA).

All the ligation products were subsequently amplified by PCR using U1cas-up ForNheI and U1cas-down RevNheI primers. The PCR reaction was performed according to the manufacturer’s protocol of Cloned *Pfu* DNA polymerase Taq (Agilent Technology, Santa Clara, CA, USA).

After amplification and purification, the resulting fragments and ten μg of the pAAV-2.1CMVeGFP vector were digested with NheI restriction enzyme (New England BioLabs, NEB, Ipswich, MA) in 50 μl final volume. The PCR products were then cloned in the forward orientation into a pAAV-2.1CMVeGFP plasmid at the NheI restriction site upstream of the CMV promoter.^55^ Ligation was performed as described above, using T4 DNA ligase, (New England BioLabs, NEB, Ipswich, MA). We confirmed the exact sequence and the orientation of the inserts within the pAAV-2.1CMVeGFP vector by automated DNA sequencing (BMR Genomics, Padova), using the AAV Rev primer (5’-ccatatatgggctatgaataatg-3’).

### Binding energy (ΔG°_37_) prediction

RNAfold algorithms were used to predict secondary structures and the minimum free energy (ΔG°_37_, kcal/mol) for the RPGR pre-mRNA^56,57^ (http://rna.tbi.univie.ac.at//cgi-bin/RNAWebSuite/RNAfold.cgi). The DuplexFold, from the package RNAstructure 3.5^58^ (http://rna.urmc.rochester.edu/RNAstructureWeb/), was used to calculate the binding energy (ΔG°_37_ overall, kcal/mol) for each RPGR mRNA-Chimeric RNA interaction.

### Cell culture and transfection conditions

Wild-type and mutant RPGR minigenes (MINI wt, MINI mut) and chimeric U1_snRNA constructs (U1_3’, U1_5’, U1_3’5’ and U1_Scramble) were transiently co-transfected into Human embryonic kidney (HEK-293T) and rat Pheochromocytoma (PC-12) cell lines. HEK-293T cells were maintained in Dulbecco’s Modified Eagle’s Medium (DMEM) with red phenol supplemented with 10% Fetal Bovine Serum (FBS), 2mM Glutamine, 100 U/μl Penicillin/Streptomycin, and were grown in a humidified incubator at 37°C and 5% CO2. Lipofectamine 2000 (Life-Technologies, Carlsbad, CA) was used for co-transfection with RPGR minigenes and chimeric U1 snRNAs, according to the manufacturer’s instructions.

Rat PC-12 cells (ATCC entry CRL-1721) were grown at 37°C (5 % CO2) in supplemented DMEM: Dulbecco’s Modified Eagle’s Medium with 4.5 % glucose (Lonza, Visp, Switzerland) supplemented with 10 % fetal bovine serum (Gibco, Grand Island, NY, USA), 5 % horse serum (Gibco), 1 mM glutamine (Gibco), 1 mM Penicillin/Streptomycin (Gibco). Cells were seeded in T-75 cm^2^ flasks (Corning, NY, USA) coated with 50 ng/ml poly-D-lysine hydrobromide (Sigma, St. Louis, MO, USA), to achieve 70 % confluence.

The same amount of each U1_asRNA plasmids (200 ng) was used either alone (U1_3’, U1_5’, U1_3’5’ or U1_Scramble) or in combination (U1_3’(100 ng) + U1_5’(100ng)). For each co-transfection experiment, the same amount of MINI mut or MINI wt was used (400 ng).

Transfection of PC-12 cells with RPGR minigenes and chimeric U1 snRNAs was performed using the Neon-Transfection System MPK5000 (Life-Technologies, Carlsbad, CA) under the following conditions: three 10.ms long pulses each one of 1500V as described in Covello et al. (2014).^37^ HEK-293T (4.0×10^4^ cells/well) and PC-12 (3.0×10^5^ cells/well) cells were plated onto 24-multi-well plates. An amount of 400 ng of each RPGR minigenes (MINI wt or MINI mut) plus 200 ng of each different chimeric U1 snRNAs was used for co-transfection of both HEK-293T and PC-12 cells unless specifically mentioned. After 48 hours of co-transfection, cells were trypsinized and subsequently washed once with phosphate-buffered saline (PBS). Pellets were collected and stored at −80°C for RNA analysis.

### RNA extraction and semiquantitative RT-PCR analyses

Total RNA was extracted from HEK-293T, and PC-12 cells, using Trizol Reagent (Life Technologies, Carlsbad, CA) according to the manufacturer’s instructions. RNA were treated with DNase (TURBO DNA-free Kit, Life Technologies, Carlsbad, CA, USA) and concentrations checked by NanoDrop ND-1000 Spectrophotometer (Nanodrop Technologies, Wilmington, USA). cDNA synthesis was performed using random hexamer primers and RevertAid First Strand cDNA Synthesis Kit, according to the manufacturer’s protocols (Thermo Scientific, Waltham, MA).

For all semiquantitative RT-PCR analyses, we applied the following amplification protocol, a one minute denaturation at 94°C, 35 amplification cycles (30 seconds at 94°C, 30 seconds at 58°C, and 1 minute at 72°C), and a final extension for 7 minutes at 72°C. PCR reactions were carried out in 25 μl final volume in a reaction mixture containing 1 U Taq DNA polymerase (Applied Biosystems by Life Technologies, Carlsbad, CA, USA), according to manufacturer’s protocols.

The endogenous RPGR expression levels in both cell lines HEK-293T and PC-12 were detected by semiquantitative RT-PCR assay, by using two pairs of specific primers:

human RPGR: hRPGR_E7_For: 5’-caatcacagaacaccccaga-3’; hRPGR_E10_Rev: 5’-tgacatccaccacaagcaacc-3’

rat RPGR: rRPGR_E7_For: 5’ tgatcaatcacagatctcc-3’; rRPGR_E10_Rev: 5’-tgacatccaccacaggcaatc-3’.

These primers discriminate between the two alternative endogenous RPGR transcripts (E9a+ and E9a-) producing two different amplicons: 118 bp for the isoform without E9a (E9a-) and 324 bp for the isoform with E9a (E9a+).

To analyse the transcripts deriving from wild-type and mutant minigene reporters, the semiquantitative RT-PCR assay of cDNAs from HEK-293T and PC-12 cells was performed using primers: RPGRE9-For; 5’-cggccttatgtatacttttgg-3’ and BGH-Rev; 5’-tagaaggcacagtcgagg-3’ that anneal to the RPGR exons 9 and BGH plasmid region, respectively.

The pair of primers produce an amplicon of 392 bp for the isoform without E9a (E9a-) and an amplicon of 528 bp for the isoform with E9a (E9a+). An amplicon of 496 bp from GAPDH region was amplified using primers GAPDH For; 5’-tgacctcaactacatggtctaca-3’ and GAPDH Rev; 5’-cttcccattctcggccttg-3’ and was used as an internal control (house-keeping gene). Densitometric analyses were carried out with the Imagelab 2.0 software (Biorad, Hercules, CA). After background correction, bands intensities were normalized to the GAPDH levels. The assay was performed in triplicate and the mean ± SD was calculated.

### RT-PCR assay to evaluate U1_asRNA expression in co-transfected cells

The expression levels and the integrity of the U1 snRNA chimeric constructs transfected in HEK-293T and PC-12 cells were analysed via semiquantitative RT-PCR. The primer U1+130_Rev; 5’-agcacatccggagtgcaatg-3’, which anneals to the body of the U1 mRNA molecule,^54^ was used in combination with a specific primer for the engineered antisense modified U1 tail: U1-RPGR3’_For: 5’-ccaccgtgccctgccaaaacg-3’; U1-RPGR5’_For: 5’-gccatcatacctggctaatttttg-3’ or U1-Scramble_For: 5’-gctgcgtctcgcggaataattga-3’. To analyse the expression level of double antisense U1_3’5’ construct the U1+130 Rev and U1-RPGR5’_For primers were used. PCR reactions were performed as described above for semiquantitative RT-PCR with an annealing temperature of 54°C. The U1_asRNA amplification products had a size of approximately 100 bp.

### RNA Immunoprecipitation (RIP) assay

HEK-293T cells were grown in 150 mm culture dishes (1.8 × 10^7^ cells/dish) at 37°C in 5% CO2 atmosphere. After 24 hours, the cells were co-transfected with U1 snRNAs constructs (U1_3’, U1_5’ and U1_Scramble) (c.a. 15 μg) and RPGR mutant minigene (MINI mut) (c.a. 28 μg) using the Lipofectamine 2000 method according to the manufacturer’s protocol (Life-Technologies, Carlsbad, CA). 48 hours after co-transfection, the culture medium was removed, and ice-cold phosphate-buffered saline (PBS) was added to the cells. The culture dishes were placed on ice and irradiated once with 150 mJ/cm2 at 350 UV for 1 minute using Ultraviolet Crosslinker, UVP CL-1000L Model: 365nm UV (Thermo Fisher Scientific, Illikirch Cedex, France).

Cells were harvested with a cell scraper, centrifuged briefly to remove PBS, and then nuclear proteins fraction was obtained from HEK-293T cells by a modified version of the protocol described by Dignam and collaborators.^59^ Briefly, the pellet was harvested, resuspended into 5X Cytoplasmic Extract (CE) buffer with NP-40 (10 mM HEPES pH 7.0, 10 mM KCl, 1.5 mM MgCl2, 0.1 M EDTA, 0.6% (v/v) NP-40, 1mM DTT, 100U/ml −1 RNase out (Life-Technologies, Carlsbad, CA), 2 U/ml Turbo Dnase (Ambion, Life-Technologies, Carlsbad, CA) and proteinase inhibitor cocktail (Sigma Aldrich, St. Louis, MO)), adjusted to pH 7.6. After centrifugation (1,200 rpm for 10 minutes at 4°C), cells were resuspended in CE buffer without NP-40. Cells were centrifuged at 4°C (3,000 rpm for 5 minutes), and an equal volume of Nuclear Extract (NE) buffer (20 mM HEPES pH7.9, 400 mM NaCl, 1 mM EDTA, 1X Proteinase inhibitor cocktail (Sigma Aldrich, St. Louis, MO), 100U/ml-1 RNase out (Life-Technologies, Carlsbad, CA), 2 U/ml Turbo DNase (Ambion, Life-Technologies, Carlsbad, CA), and 25% (v/v) glycerol, adjusted to pH 8.0) was added to this pellet and was kept on ice for 10 min. After homogenization, the cell suspension was centrifuged for 5 minutes at 14,000 rpm at 4°C. Final proteins concentration in the NE extracts was determined using the colorimetric Pierce BCA protein assay kit (Thermo Scientific, Waltham, MA). Pellets were stored at −80° C.

Immunoprecipitation was carried out from the nuclear fraction by Dynabeads protein A (Life Technologies, Carlsbad, CA) with the Anti-U1A antibody (U1-70K) (Abcam, Cambridge, UK) and the Normal Rabbit IgG (Millipore, Darmstadt, Germany) as a negative control. Five μl of both antibodies (anti-U1-70k and IgG) were used with 200 μl of the nuclear extract, and the Rip-Chip assay was performed as described by Keene, et al. (2006)^38^ that was optimized to minimize inappropriate interaction.

RNAs were extracted from immunoprecipitation fractions using Trizol reagent, according to the manufacturer’s instructions (Trizol Reagent, Life Technologies, Carlsbad, CA) and were treated with DNase (TURBO DNA-free Kit, Life Technologies, Carlsbad, CA, USA). RNA was used to perform U1_snRNA chimeric screening, according to the procedure reported in the section “RT-PCR assay to evaluate U1_asRNA expression in co-transfected cells. 10 μl PCR products were analysed on 2.5% agarose gel. The amplicons were about 100 bp long, as predicted.

### Statistical analyses

Semiquantitative RT-PCR densitometric data were expressed as means ± SD (n=3) and were compared using two-way ANOVA, followed by Bonferroni’s comparisons test, for multiple comparisons test, and unpaired two-tailed Student t-test, for two groups comparison (GraphPad Software, San Diego, CA. www.graphpad.com). Statistical significance is denoted with asterisks (*P<0.05; **P<0.01; ***P<0.001; ****P<0.0001).

## Supplemental information

Supplemental information includes one table and five figures can be found at the end of this article.

## Author contributions

M.A.D. and S.C. conceived the study, designed the experiments, interpreted the results and critically reviewed the manuscript; M.A.D. and G.C. wrote the paper; G.C. contributed to the experimental design, performed the experiments, analysed the data and prepared the figures; G.H. and N.B. conducted the experiments and performed bibliographic researches. All authors reviewed the manuscript.

## Acknowledgements

This work was funded by the Italian Ministry of Health (Project GR-2008-1136933), by Department CIBIO, University of Trento (Grant number 40201033) to M.A.D., and supported by COST Actions-BM1207 - Networking towards clinical application of antisense-mediated exon skipping and by COST Action CA17103 - Delivery of Antisense RNA Therapeutics. We would like to acknowledge Annalisa Rossi (Laboratory of Transcriptional Networks, Department CIBIO, University of Trento) for the helpful suggestions in regard to RIP assay protocol.

## Supplemental material

**Supplemental Figure S1.**
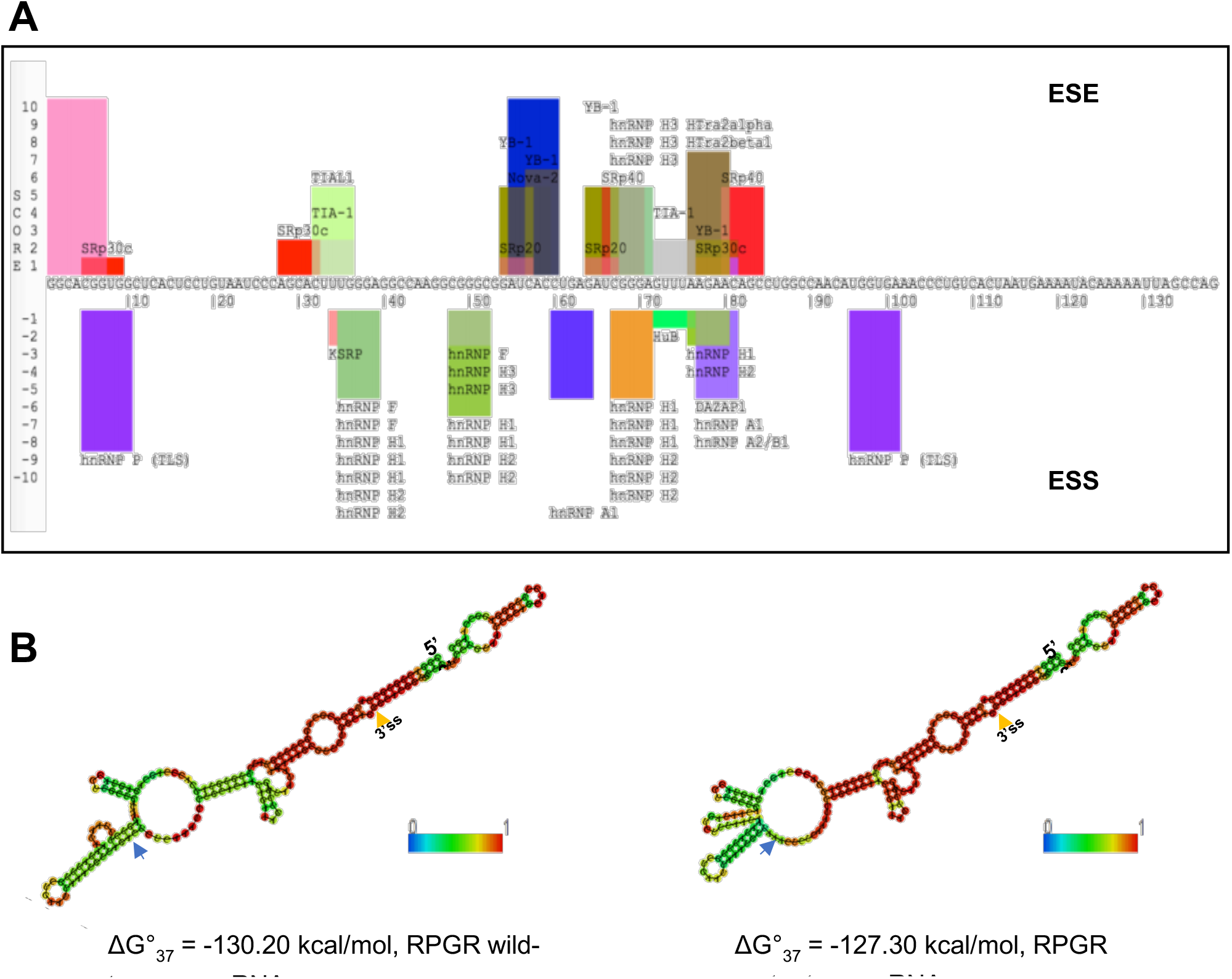
Computational analysis of splicing sequences in RPGR exon 9a. **(a)** Prediction for Exonic Splice Enhances (ESE) and Exonic Splice Silencers (ESS) in RPGR exon 9a using SpliceAid. All predicted ESEs are overlapping with predicted ESS thus a clear ESE is not defined. **(b)** Folding of a portion of the wild-type and mutant RPGR pre-mRNA sequence (from position 26526 to position 26728 of NCBI Refseq: NG_009553.1) and ΔG°_37_ free energy was predicted by using RNAfold algorithms. Blue arrows indicate the nucleotide position G and A in wild-type or mutant pre-mRNA sequences of RPGR. The 3’ss of exon 9a is indicated by a yellow arrow. Colour code bar (0 to 1) indicates base-pairing probability.

**Supplemental Figure S2.**
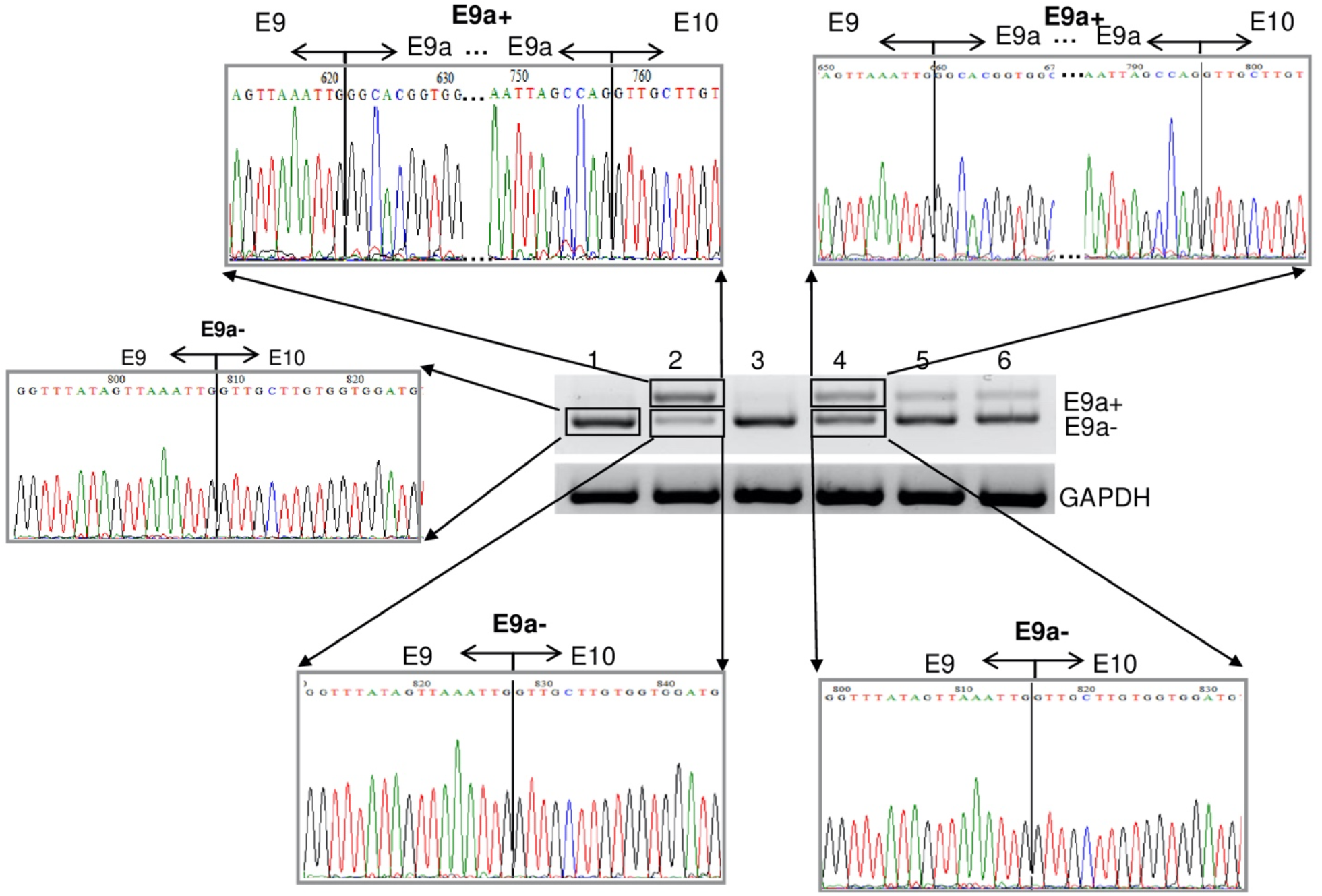
Sequencing of amplicons in HEK-293T transfections. The E9a- band from lane 1 and both the E9a- and the E9a+ bands from lanes 2 and 4 of the gel in Figure 4A, were eluted and sequenced. The chromatograms indicate a correct fusion between exons 9 and 10 for the E9a- bands and that the upper E9a+ bands correspond to the retained exon 9a.

**Supplemental Figure S3.**
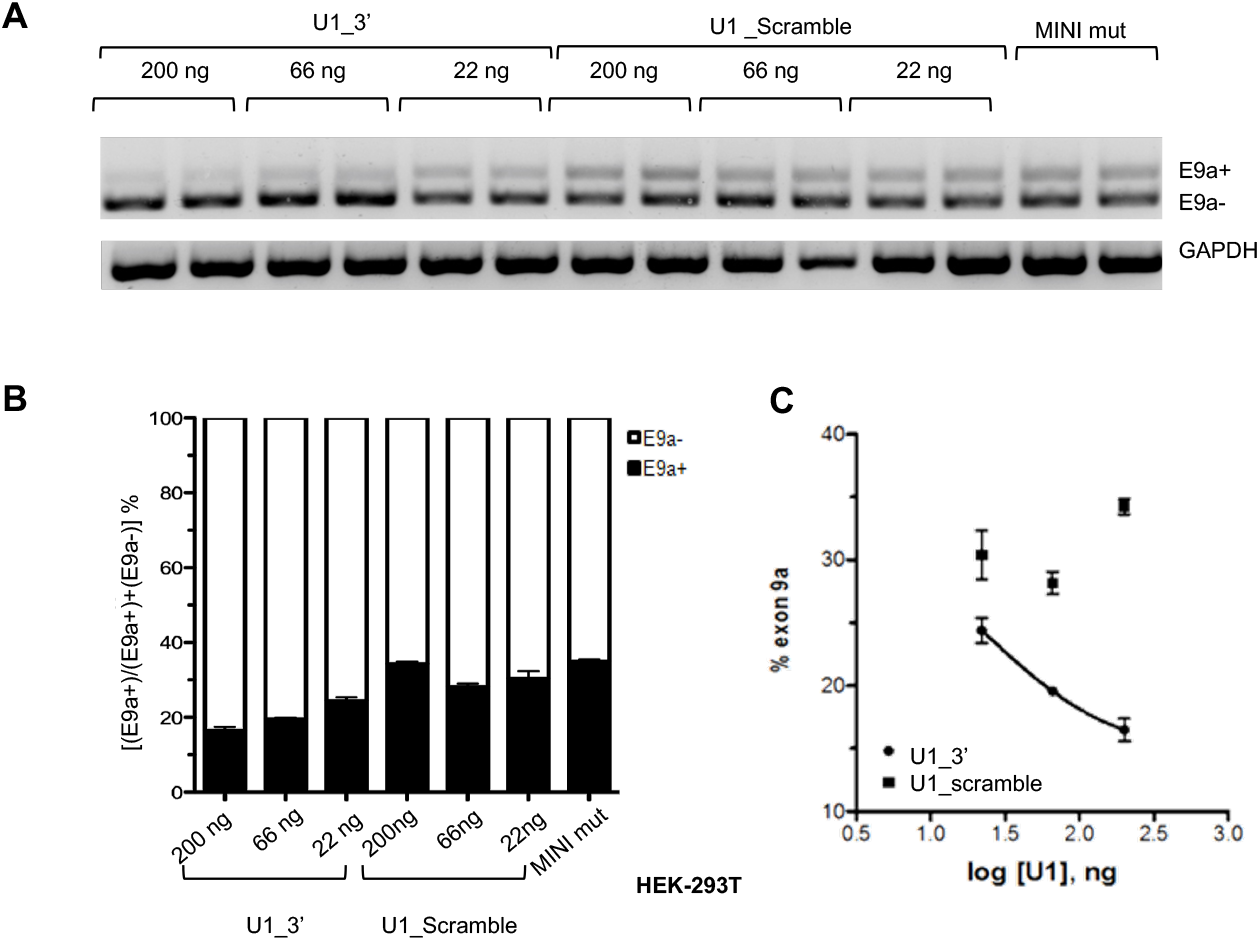
Dose-dependent skipping effect of U1_3’ and U1_Scramble. **(a)** Semiquantitative RT-PCR of RPGR minigene transcripts in HEK-293T cells co-transfected with the RPGR mutant minigene with different amounts of U1_3’ and U1_Scramble plasmids (200, 66 and 20 ng). For each plasmid concentration, two biological replicates were analysed and are shown. **(b)** Densitometric analyses of the bands (n=2). **(c)** Dose-response curve of U1_3’ and U1_Scramble. Only data of U1_3’ dilutions fit a dose-response curve.

**Supplemental Figure S4.**
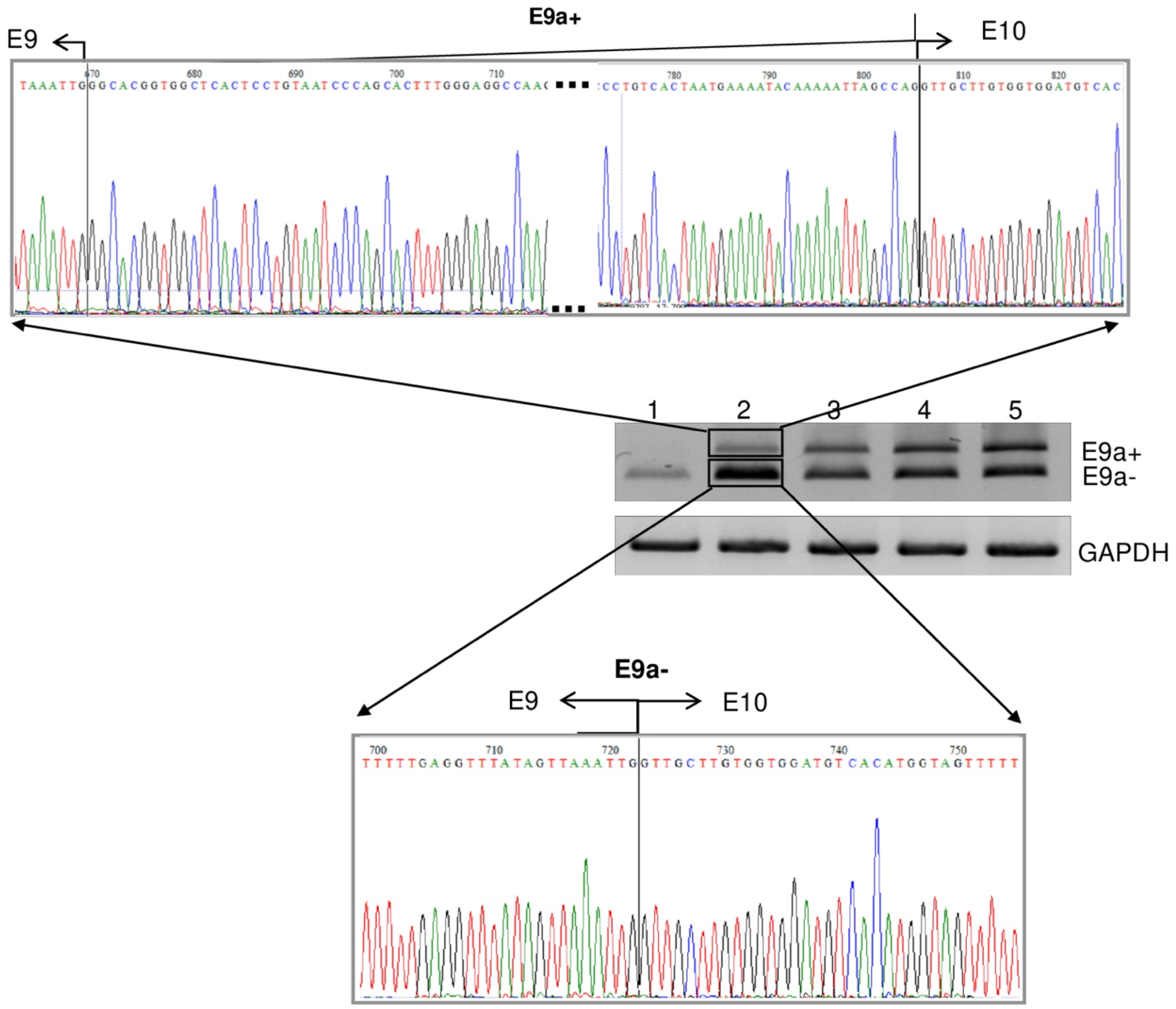
Sequencing of amplicons in PC-12 transfections. The E9a- and the E9a+ bands from lanes 2 of the gel in Figure 4B, were eluted and sequenced. The chromatograms indicate that a correct fusion between exons 9 and 10 for the E9a- bands and that the upper E9a+ bands correspond to the retained exon 9a.

**Supplemental Figure S5.**
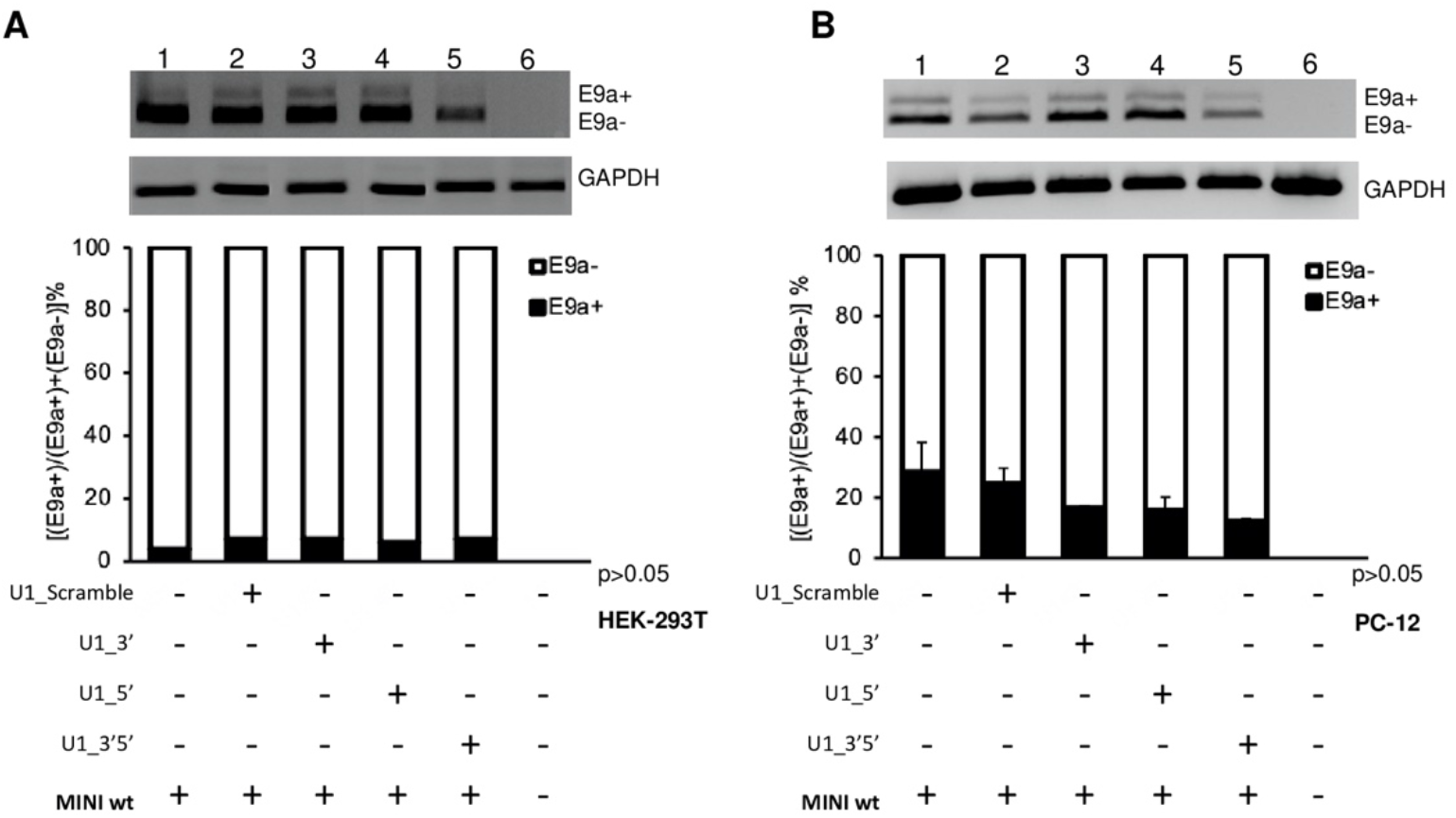
Co-transfection of RPGR MINI wt with U1 chimeric constructs. **(a)** Semiquantitative RT-PCR of RNA from HEK-293T cells transfected with RPGR wild-type minigene (MIN wt) alone or in combination with chimeric U1_snRNAs plasmids. **(b)** Semiquantitative RT-PCR of RNA from PC-12 cells transfected with RPGR wild-type minigene (MINI wt) alone or in combination with chimeric U1 snRNAs. One representative gel of three is shown in both (a) and (b). Densitometric analysis of E9a+ and E9a- amplicons, from three independent experiments, is shown for both cell lines. GAPDH is used as an internal control. Data are shown as mean ± S.D (n=3).

**Supplemental Table S.1.**
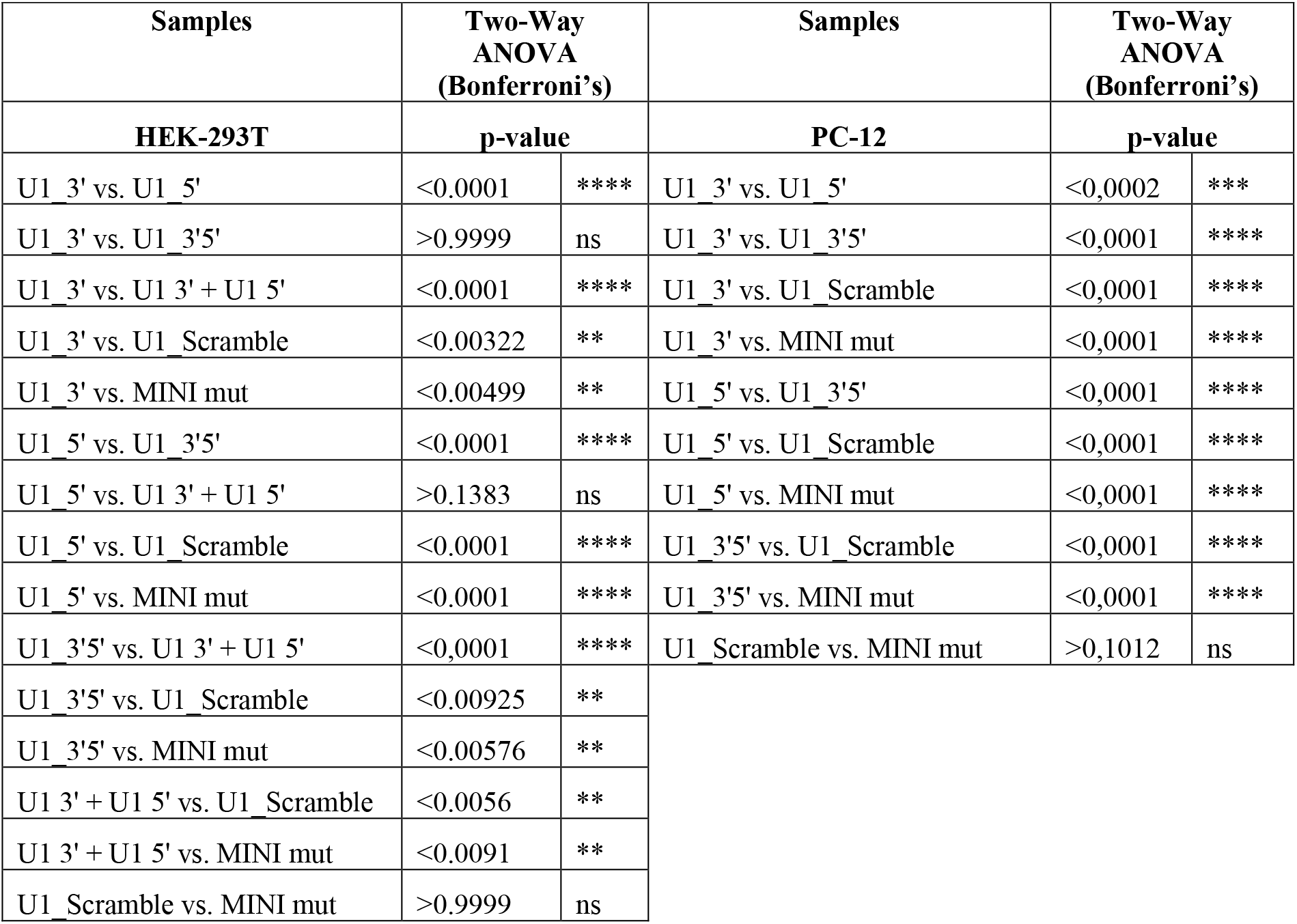
A global statistical test (GST) analysis carried out in HEK293-T and PC-12 semiquantitative RT-PCR data. p-value: *P < 0.05; **P < 0.01; ***P < 0.001 ****P < 0.0001; ns: not significant

